# Enzyme Aggregation and Fragmentation Induced by Catalysis Relevant Species

**DOI:** 10.1101/2021.09.03.458882

**Authors:** Kayla Gentile, Ashlesha Bhide, Joshua Kauffman, Subhadip Ghosh, Subhabrata Maiti, James Adair, Tae Hee Lee, Ayusman Sen

## Abstract

It is usually assumed that enzymes retain their native structure during catalysis. However, the aggregation and fragmentation of proteins can be difficult to detect and sometimes conclusions are drawn based on the assumption that the protein is in its native form. We have examined three model enzymes, alkaline phosphatase (AkP), hexokinase (HK) and glucose oxidase (GOx). We find that these enzymes aggregate or fragment after addition of chemical species directly related to their catalysis. We used several independent techniques to study this behavior. Specifically, we found that glucose oxidase and hexokinase fragment in the presence of D-Glucose but not L-glucose, while hexokinase aggregates in the presence of Mg^2+^ ion and either ATP or ADP at low pH. Alkaline phosphatase aggregates in the presence of Zn^2+^ ion and inorganic phosphate. The aggregation of hexokinase and alkaline phosphatase does not appear to attenuate their catalytic activity. Our study indicates that specific multimeric structures of native enzymes may not be retained during catalysis and suggests pathways for different enzymes to associate or separate over the course of substrate turnover.

## Introduction

Enzyme catalysis is critical to the viability of living systems.^1^ Enzymes are also employed increasingly in a myriad of technological applications.^2–4^ Generally, it is assumed that enzymes retain their native structure during catalysis. However, the aggregation and fragmentation of proteins can be difficult to detect and sometimes conclusions are drawn based on the assumption that the protein is in its native form.^5^ This can be particularly problematical during enzyme purification and immobilization.^6^

Many enzymes have been shown to exhibit enhanced diffusion while catalyzing the turnover of their substrates.^7–16^ However, there are hints that experimental artifacts may be vitiating some of the observations.^17–19^ One suggested possibility is that some enzymes may be fragmenting into their monomeric units during catalysis.^8,20^ Due to the Stokes-Einstein relationship, size is inversely correlated to diffusion and so smaller sized particles will diffuse faster than larger particles. Thus, it is essential to understand how the size of the enzyme particles change in the course of catalysis.

In this study, we evaluated three model enzymes, alkaline phosphatase (AkP), hexokinase (HK), and glucose oxidase (GOx). We find that these enzymes either aggregate or fragment following the addition of simple chemicals that are relevant for their catalytic activity. We use several independent techniques to study this behavior: fluorescence resonance energy transfer (FRET) and dynamic light scattering (DLS), along with liquid atomic force microscopy (AFM). Understanding the aggregation/fragmentation of the enzymes is critical to elucidating their functional properties.

## Materials and Methods

### Fluorescent labeling of AkP, HK, GOx

Alkaline phosphatase (from bovine intestinal mucosa), hexokinase (from Saccharomyces cerevisiae), glucose oxidase (from *Aspergillus niger*), and invertase (from baker’s yeast (*S. cerevisiae*) were all purchased from Sigma-Aldrich-Millipore. For experiments with tagged enzymes, each enzyme was divided into two populations and each population was tagged with an amine reactive dye, either Alexa Fluor 488 (AF488; ex/em: 490/525; Thermo Fisher Scientific) or Alexa Fluor 532 (AF532; ex/em: 532/554; Thermo Fisher Scientific). The emission spectra of AF488 and AF532 are shown in **Figure S1**.

For example, alkaline phosphatase (12.5 µM) was reacted with a six-fold excess of Alexa Fluor 488 and alkaline phosphatase (12.5 µM) was reacted with a two-fold excess of Alexa Fluor 532 in water, along with 100 mM sodium bicarbonate, for 1 hour on a rotator. Then the mixtures were allowed to sit in the 4ºC fridge overnight and were allowed to rotate again the next morning for about 2 hours. The enzyme-dye mixtures were purified according to protocol included with the Antibody Conjugate Purification Kit for 0.5-1 mg (Thermo Fisher Scientific). The buffer was replaced with 50 mM HEPES (pH 7; Sigma-Aldrich) for alkaline phosphatase and hexokinase and 50 mM MES (pH 6; Sigma-Aldrich) for glucose oxidase and invertase. The same procedure was followed for all enzymes with varying starting concentrations and enzyme:dye ratios. These are specified in the Supporting Information (SI) in **Table S1**.

### Fluorescence Resonance Energy Transfer (FRET) Measurements

FRET measurements were taken on a Fluorolog JobinYvon Horiba spectrofluorometer. The slit width was set to 5, the integration time to 0.5 s, the increment to 5 nm and the detector to S1/R1. Spectra were recorded on FluorEssence software and analyzed using OriginPro.

#### Alkaline Phosphatase FRET Measurements

For the FRET experiments, a Micro Hellma^®^ fluorescence cuvette was used. For alkaline phosphatase, an enzyme mixture was made with 0.1 µM AkP-488 and 0.1 µM AkP-532 in 50 mM HEPES (pH 7; Sigma-Aldrich) buffer. For the zinc nitrate and magnesium nitrate titrations, zinc nitrate hexahydrate (Zn(NO_3_)_2_; Sigma-Aldrich) or magnesium nitrate hexahydrate (Mg(NO_3_)_2_; Sigma-Aldrich) was added so that the final concentration of salt for each experiment was 0, 0.05, 0.1, 0.5, 1, 2.5, 5, 7.5, or 10 mM. Fluorescence was recorded with just the enzyme before salt was added. The fluorescence reading with salt was taken 1 minute after the addition of salt. The Hofmeister salt experiments were conducted in a similar manner. The salts used were ammonium nitrate (NH_4_NO_3_; Sigma-Aldrich), sodium nitrate (NaNO_3_; Alfa Aesar), calcium nitrate tetrahydrate (Ca(NO_3_)_2_; Sigma-Aldrich), sodium sulfate anhydrous (Na_2_SO_4_; BDH), sodium phosphate dibasic hexahydrate (Na_2_HPO_4_; Sigma-Aldrich), sodium chloride (NaCl; Sigma-Aldrich) and sodium thiocyanate (NaSCN; Sigma-Aldrich). The salt was added so that the final concentration was 1 mM and fluorescence readings were taken with just the enzyme and then 1 minute after salt was added.

For the alkaline phosphatase reaction experiments, zinc nitrate hexahydrate (Zn(NO_3_)_2_; Sigma-Aldrich) or magnesium nitrate hexahydrate (Mg(NO_3_)_2_; Sigma-Aldrich), D-Glucose 6-phosphate sodium salt (Sigma-Aldrich), D-(+)-Glucose (Sigma-Aldrich) or sodium phosphate dibasic hexahydrate (Sigma-Aldrich) were added to the experimental solution. All substrates were added so that the final concentration for each substrate was 0.5 mM. Ethylenediamine tetraacetic acid disodium salt (EDTA; IBI Scientific) was added in the last 5 minutes to experiments so that its final concentration was 0.5 mM. For the alkaline phosphatase fragmentation experiments D-(+)-Glucose (D-Glu; Sigma Aldrich, 20 mM) or L-(–)-Glucose (L-Glu; Sigma Aldrich, 20 mM) were used.

#### Hexokinase FRET Measurements

For the experiments, a Micro Hellma^®^ fluorescence cuvette was used. For hexokinase, an enzyme mixture was made with 0.1 µM HK-488 and 0.1 µM HK-532 in 50 mM HEPES (pH 7; Sigma-Aldrich) buffer. The titration and Hofmeister salt experiments were executed in the same manner as the alkaline phosphatase experiments with the same reagents, but a solution of 0.1 µM HK-488 and 0.1 µM HK-532 was used. In addition, magnesium chloride anhydrous (MgCl_2_; Alfa Aesar) was used for the titration instead of zinc nitrate or magnesium nitrate. For the hexokinase reaction experiments, magnesium chloride anhydrous (MgCl_2_; Alfa Aesar) was added to all experiments so that the final concentration was 40 mM. Adenosine 5’-triphosphate disodium salt hydrate (ATP; Sigma-Aldrich), D-(+)-Glucose (Glu; Sigma-Aldrich), D-Glucose 6-phosphate sodium salt (G6P; Sigma-Aldrich), or Adenosine 5’-diphosphate sodium salt (ADP; Sigma-Aldrich) was added so the final concentration for each substrate was 20 mM.

For the pH experiments described in **Figure 7**, two solutions of 500 mM ATP and two solutions of 500 mM ADP were made in 50 mM HEPES (pH 7; Sigma-Aldrich). The pH was measured using a Thermo Scientific Orion Star pH meter. For one solution of ATP and one solution of ADP, the pH was adjusted to pH 7 using 3M sodium hydroxide (Alfa Aesar), a summary of the resulting pH values can be found in **Table S2**. Then approximately 30 µL of the 500 mM ATP or ADP stock solutions were added to the experimental FRET solutions to achieve a final concentration of 20 mM ATP or ADP. **Table S3** gives the resulting pH values for the experiment recorded in **Figure 7**. For the hexokinase fragmentation experiments, D-(+)-Glucose (D-Glu; Sigma-Aldrich, 20 mM) or L-(–)-Glucose (L-Glu; Sigma-Aldrich, 20 mM) were used.

#### Glucose Oxidase FRET Measurements

For the experiments, a Micro Hellma^®^ fluorescence cuvette was used. For glucose oxidase, an enzyme mixture was made with 0.1 µM GOx-488 and 0.1 µM GOx-532 in a 50 mM MES (pH 6; Sigma-Aldrich) buffer solution. D-(+)-Glucose (D-Glu; Sigma-Aldrich), L-(-)-Glucose (L-Glu; Sigma-Aldrich), D-(+)-Gluconic acid *δ*-lactone (Sigma-Aldrich), or hydrogen peroxide, 30% (VWR) were added so that the final concentration for each substrate was 1 mM (**Figure 10**). For the experiment with invertase (**Figure 12**), a stock solution was made with 0.1 µM GOx-488 and 0.1 µM GOx-532 in a 50 mM MES (pH 6) buffer solution. Invertase was added so that the final concentration was 0.1 µM. Sucrose (Sigma-Aldrich) was added to the solution so that the final concentration was 1 mM. For determining the concentration of D-Glucose required for fragmentation (**Figure S8**), D-(+)-Glucose (D-Glu; Sigma-Aldrich) in the required quantity was added to the stock solution of 0.1 µM GOx-488 and 0.1 µM GOx-532 so that the final concentration was as specified.

### Dynamic Light Scattering (DLS) Measurements

The enzymes were analyzed on a NanoBrook Omni instrument. To reduce dust and debris that affect sample measurements, enzymes and buffer were centrifuged twice in 300kDa centrifugal filters at 3000 xg for 20 minutes using a Thermo Scientific Sorvall ST16 Centrifuge. Additionally, all enzyme samples were filtered twice with a 0.2 µm cellulose acetate syringe filter (VWR) immediately before running the experiment. Lastly, all pipette tips, cuvettes (plastic disposable; 4.5 mL) and syringes were rinsed with DI water prior to experimentation. The following settings were used for the DLS measurements: angle: 90°; correlator layout: proteins; cell type: BI-SCP; set duration: 120 sec; equilibration times: 180s; dust filter: 10-50 nm; liquid: water; baseline normalization: auto (slope analysis); threshold: 2.0 nm-5000 nm. Series of 3 or 6 runs were used to obtain data. The same reagents from the FRET experiments were used to conduct the DLS experiments with 3 mL total sample volumes, but at higher concentrations. These are specified in **Figures 4, S3 and S7**. The DLS is not sensitive enough to detect the enzymes at the low concentrations used for FRET.

### Atomic Force Microscopy (AFM) Measurements

A Bruker BioScope Resolve AFM and Bruker ScanAsyst-Fluid+ probe was used to obtain control and experimental data. Scan rate was set at 0.501 Hz with 128 samples per line and a drive amplitude of 100 mV. For control experiments, 0.2 µM alkaline phosphatase in HEPES buffer (pH 7; Sigma-Aldrich) was added to a mica surface and visualized. Zn(NO_3_)_2_ (0.5 mM; Sigma-Aldrich), Mg(NO_3_)_2_ (0.5 mM; Sigma-Aldrich), and Na_2_HPO_4_ (0.5 mM; Sigma-Aldrich) were then added to a solution of 0.2 µM alkaline phosphatase and aggregation was characterized. Substrate and buffer solutions were filtered once with a 0.2 µM cellulose acetate syringe filter (VWR) before being added to the mica.

### Enzyme Activity Measurements

#### Alkaline Phosphatase Activity Measurements

Alkaline phosphatase was tagged according to procedure stated in the fluorescent tagging section. Activity was monitored by measuring absorbance using a Thermo Scientific Evolution 220 UV-Visible Spectrophotometer. For alkaline phosphatase, the reaction illustrated in Equation 2 was used. The reaction product, para-nitro phenol (pNP), is UV active at 405 nm and the activity can be monitored by following the production of pNP over time. An assay mixture, 1 mL in total volume, contained 0.2 µM tagged AkP, 1 mM p-Nitrophenyl phosphate disodium salt hexahydrate (pNPP; Sigma-Aldrich) and either 0, 0.1, 0.5, 1, or 5 mM of Zn(NO_3_)_2_ (Sigma-Aldrich) and Mg(NO_3_)_2_ (Sigma-Aldrich) in 50 mM HEPES (pH 7; Sigma-Aldrich).

#### Hexokinase Activity Measurements

Hexokinase was tagged according to procedure stated in the fluorescent tagging section. Activity was monitored by measuring absorbance using a Thermo Scientific Evolution 220 UV-Visible Spectrophotometer. For hexokinase, the reactions illustrated in Equations 3-4 was used. An assay mixture, 1 mL in total volume, contained 0.2 µM tagged HK, 20 mM D-(+)-Glucose, 20 mM ATP, 2.5 mM ß-Nicotinamide adenine dinucleotide phosphate sodium salt hydrate (NADP^+^; Sigma-Aldrich), 10 units of Glucose-6-phosphate Dehydrogenase from baker’s yeast (G6PDH; *S. cerevisiae*; Sigma-Aldrich) and either 0, 20 or 40 mM of MgCl_2_ in 50 mM HEPES (pH 7).

### Statistical Analysis

The statistical significance of the data sets in **Figures 2, 8, S2-S5, and S8** was evaluated using an unpaired t-test. The alpha level chosen for the t-tests was 5% (0.05). When the two-tailed P value (calculated using the unpaired t-test) was less than the alpha level (0.05), the results were described as statistically different.

## Results and Discussion

### Enzymes that Exhibit Aggregation

#### Alkaline Phosphatase aggregates with the addition of Zn^2+^ and inorganic phosphate

Alkaline phosphatase (AkP) from bovine intestinal mucosa was purchased in a lyophilized powder form from Millipore Sigma. Its structure is shown in **Figure 1**. It is a 160 kDa dimeric phosphoesterase that catalyzes the decomposition of phosphate containing compounds, including glucose 6-phosphate (G6P) and p-nitrophenyl phosphate (pNPP) (Equations 1-2).

**Fig 1:**
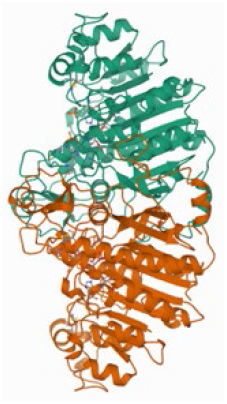
AkP Structure. Protein Data Bank (PDB) Structure of alkaline phosphatase (from *Escherichia Coli)*.^21^

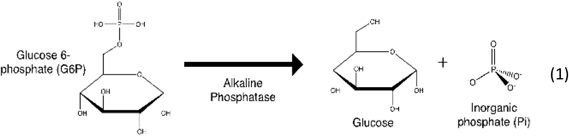

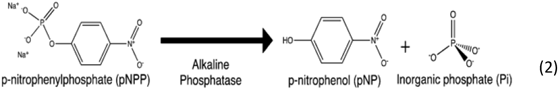

To begin, we studied the aggregation of this enzyme using fluorescence resonance energy transfer (FRET). FRET is a technique that uses two fluorophores and allows researchers to estimate how close two fluorescently tagged molecules are to each other.^22,23^ We used two fluorescent dyes in a FRET pair (AlexaFluor 488 nm and Alexa Fluor 532 nm; AF488 and AF532) to tag the enzymes and we report FRET efficiencies as a measure of the proportion of the enzyme population that is within approximately 6 nm of each other (more details can be found in the SI). The higher the FRET efficiency, the higher the proportion of enzymes that are close to each other in the experiment.

For the FRET experiments, we tagged half of the AkP population (0.1 µM) with AF488 and the other half of the population (0.1 µM) with AF532 (tagging procedures are described in the Materials and Methods). Unless otherwise noted, this is the procedure followed for all enzyme FRET studies described. Literature notes that the catalytic activity of AkP is enhanced by both zinc and magnesium ions.^24^ Thus, we carried out a FRET assay with increasing concentrations of zinc and magnesium nitrate with the tagged AkP. For reference, the free concentrations of zinc and magnesium ions in the cell are estimated to be about 0.2 – 0.3 mM and 0.5 – 1 mM, respectively.^25,26^ The results are shown in **Figure 2** and demonstrate that zinc nitrate causes a concentration dependent increase in the enzyme aggregation, while magnesium nitrate has no significant effect.

**Fig. 2.**
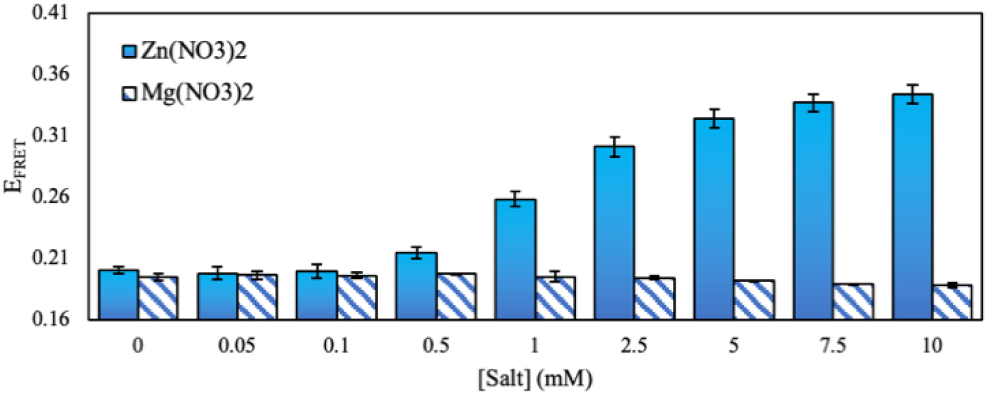
FRET efficiencies of AkP with Zn^2+^ and Mg^2+^. FRET efficiency of 0.2 µM AkP with increasing mM concentrations of Zn(NO_3_)_2_ (blue bars) and Mg(NO_3_)_2_ (striped bars). The buffer used to make all experimental solutions was 50 mM HEPES (pH 7). The error bars represent the standard deviation from two trials. The FRET efficiencies from 1-10 mM Zn(NO_3_)_2_ are statistically different from the FRET efficiency with only AkP (P < 0.05; see Materials and Methods).

We proceeded to test a variety of ions in the Hofmeister series, a series that dictates whether proteins will salt into solution (dissolve) or salt out of solution (aggregate).^27^ Surprisingly, other than the Zn^2+^ ion, these salts do not have a strong effect on enzyme aggregation, regardless of whether they are on the “salting in” or “salting out” side of the series (**Figure S2)**.

Next, we introduced the enzyme’s substrate to see if catalysis has an effect on the aggregation (**Figure 3**). First, we measured the FRET efficiency only with fluorescently tagged alkaline phosphatase (0.2 µM) to act as the baseline. For further experiments, unless otherwise noted, we added minimal zinc and magnesium ions (0.5 mM) to the enzyme solution to ensure enzyme activity. At this concentration, zinc ions will not cause significant aggregation (**Figure 2**). We examined the effect of adding the substrate glucose 6-phosphate (G6P) as well as the products of the reaction, D-Glucose (D-Glu) and inorganic phosphate (Pi; Na_2_HPO_4_). Following the addition of the substrate G6P, the FRET efficiency begins to increase after about 10 minutes and then continues to increase for the remainder of the experiment. Interestingly, an increase in FRET efficiency is observed immediately when Pi is added. However, the addition of the other product in G6P hydrolysis, D-Glu, has no effect. Therefore, we postulate that the inorganic phosphate is the cause of the aggregation of AkP. The glucose 6-phosphate reaction produces phosphate over time which is why the FRET signal increases slowly. But, if Pi is added at the start, the FRET efficiency increases immediately. It is important to note that this behavior is only observed when zinc and magnesium are present. Thus, with just alkaline phosphatase and phosphate and no Zn^2+^ or Mg^2+^ (green dashes) no aggregation is observed. Additionally, if we add the common zinc chelating agent (ethylenediaminetetraacetic acid, EDTA) to the experiment after 30 minutes, we see an immediate and significant drop in the FRET efficiency back to the baseline.

**Fig. 3.**
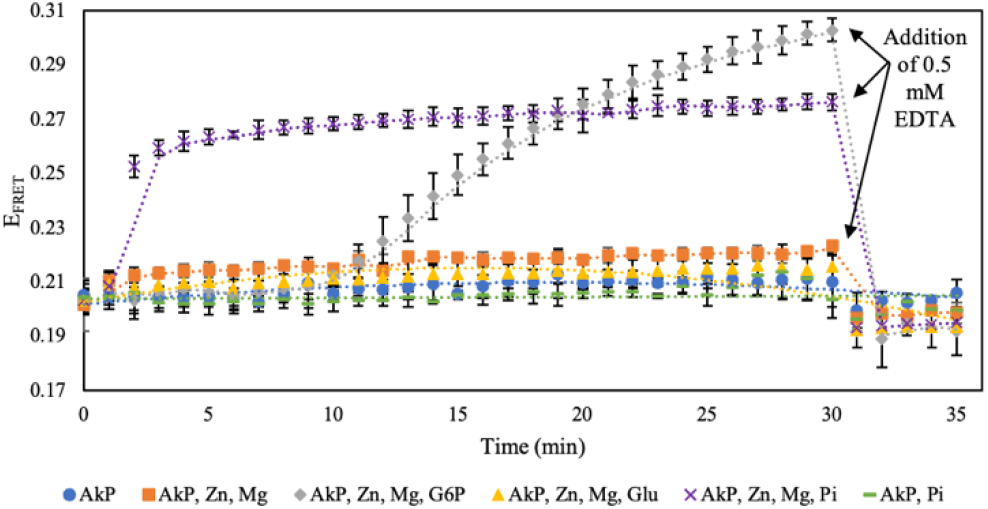
FRET efficiencies of AkP during catalysis. FRET efficiency for 0.2 µM AkP (blue circles); 0.2 µM AkP, 0.5 mM Zn(NO_3_)_2_, 0.5 mM Mg(NO_3_)_2_ (orange squares); 0.2 µM AkP, 0.5 mM Zn(NO_3_)_2_, 0.5 mM Mg(NO_3_)_2_, 0.5 mM G6P (gray diamonds); 0.2 µM AkP, 0.5 mM Zn(NO_3_)_2_, 0.5 mM Mg(NO_3_)_2_, 0.5 mM D-Glu (yellow triangles); 0.2 µM AkP, 0.5 mM Zn(NO_3_)_2_, 0.5 mM Mg(NO_3_)_2_, 0.5 mM Na_2_HPO_4_ (purple crosses); 0.2 µM AkP, 0.5 mM Na_2_HPO_4_ (green dashes). The arrows indicate the addition of 0.5 mM EDTA. The buffer used to make all experimental solutions was 50 mM HEPES (pH 7). The dashed lines are to guide the eye. The error bars represent the standard deviation from two trials.

The above results suggest that the AkP units are coming close together and aggregating. However, there is a second possibility that can give rise to higher FRET efficiencies. This involves a rapid dynamic equilibrium involving the dissociation and recombination of the enzyme subunits.^28,29^ If the subunits from the AF488 and AF532 tagged AkP molecules dissociate and recombine randomly, this will result in AkP molecules incorporating subunits tagged with both dyes, resulting in an increase in net FRET efficiency. In order to eliminate this mechanism for the observed increase in FRET efficiency and also to confirm that the fluorescent dyes or tagging procedures did not affect the results, we also examined the behavior without fluorophores by dynamic light scattering (DLS). Due to the lower sensitivity of DLS, we had to use higher concentrations of enzyme and substrate. We assessed the count rate and particle diameter versus the concentration of the aggregator or the time that the enzyme was exposed to the aggregator. Count rate is defined as the number of photons per second that the instrument detected and can be used as a measure of aggregation.^30^ Using these methods, we tested the aggregation with increasing concentrations of zinc and magnesium nitrate. Similar to the FRET results, zinc nitrate caused a concentration dependent increase in size, while magnesium nitrate had no effect (**Figure S3**).

When we added either of the two substrates for alkaline phosphatase (p-nitrophenyl phosphate, pNPP; glucose-6-phosphate, G6P), we observe a time dependent aggregation. G6P is a slower substrate than pNPP and Pi will form faster in the latter reaction.^31^ Although we observe enzyme aggregation with both substrates, the aggregation is immediate for pNPP (**Figure 4B**) but starts at around 6 minutes for G6P (**Figure 4C**), when compared to AkP without substrate (**Figure 4A**), further supporting that Pi is the aggregator. Again, the presence of zinc ions is necessary for the aggregation to occur. Overall, our findings from the DLS experiments are consistent with the FRET results.

**Fig. 4.**
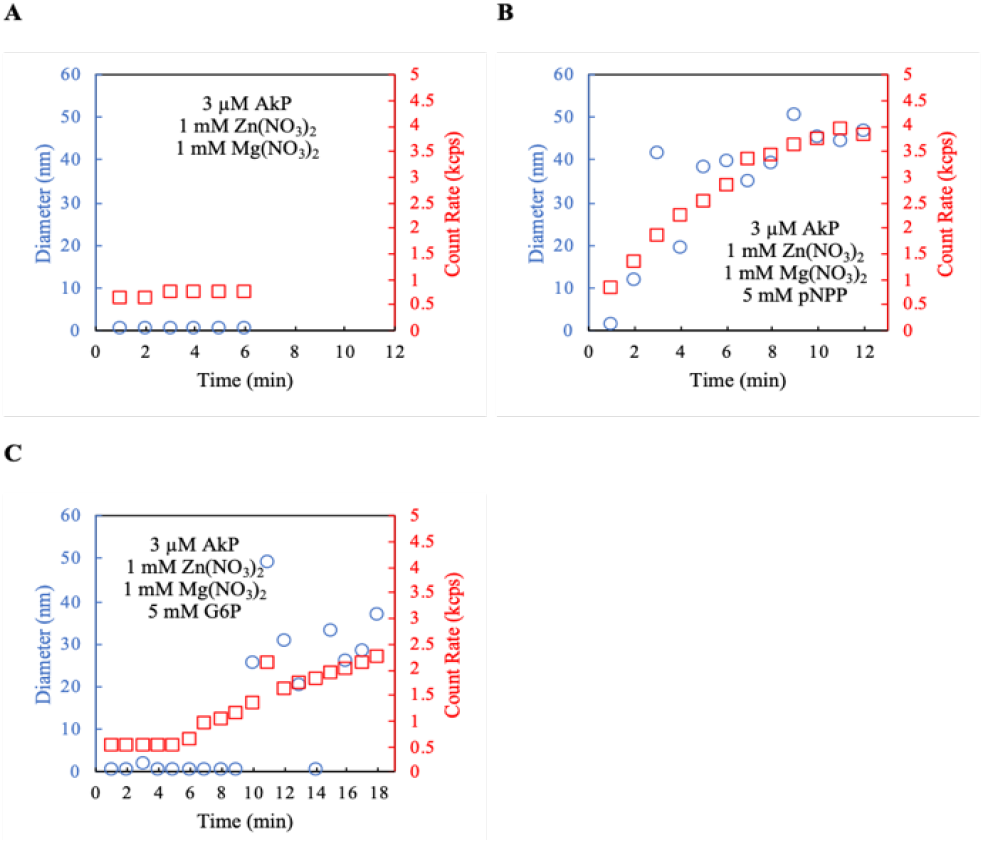
DLS data for AkP during catalysis. Diameter (blue circles) and count rate (red squares) for (A) 3 µM AkP, 1 mM Zn(NO_3_)_2_, 1 mM Mg(NO_3_)_2_ and (B) 3 µM AkP, 1 mM Zn(NO_3_)_2_, 1 mM Mg(NO_3_)_2_, 5 mM pNPP, (C) 3 µM AkP, 1 mM Zn(NO_3_)_2_, 1 mM Mg(NO_3_)_2_, mM G6P over a period of 12 to 18 minutes. The buffer used to make all experimental solutions was 50 mM HEPES (pH 7).

In addition to FRET, atomic force microscopy (AFM) was used to characterize enzyme aggregation which allows for direct visualization of the enzyme aggregation in real time. **Figure 5** shows both 2D and 3D images of alkaline phosphatase on a mica surface. Before the addition of the salts, the surface has minimal aggregates as most of the enzyme is still in solution (**Figure 5A-B**). When salts (Zn^2+^, Mg^2+^, Pi) are added, aggregates settle to the surface and can be visualized as shown in **Figure 5C-D**. Taken together, the results from the DLS and AFM experiments suggest the increase in FRET efficiency stems from the aggregation of the AkP molecules, and not from a dynamic equilibrium involving the dissociation and recombination of monomeric subunits.

**Fig. 5.**
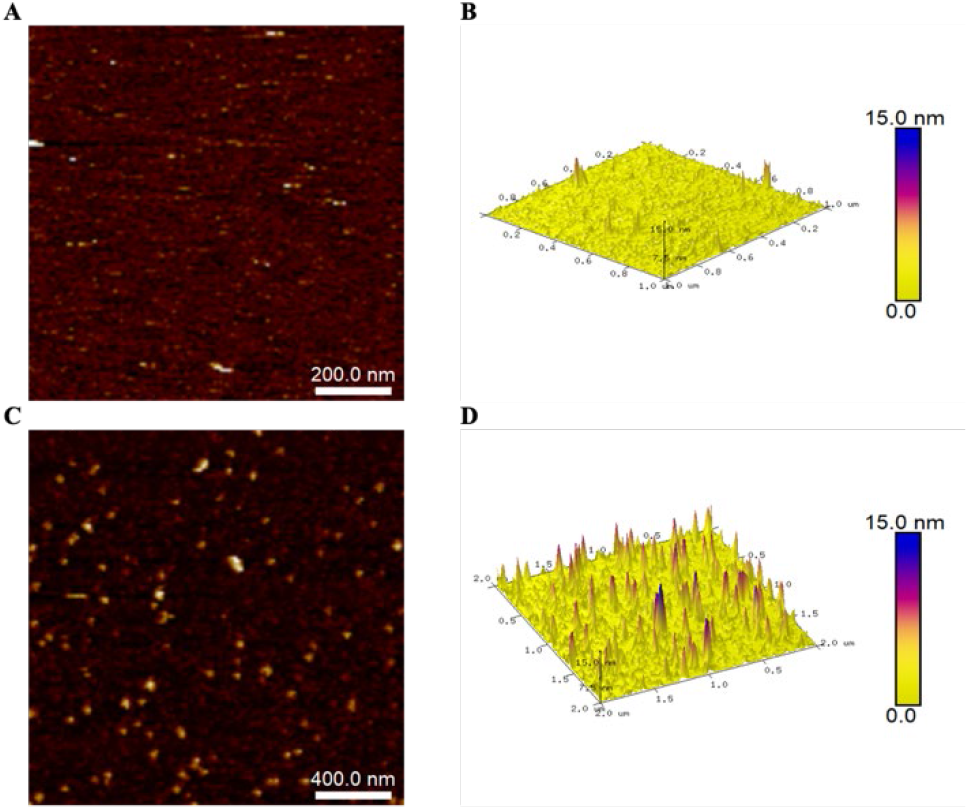
AFM data of AkP with Zn^2+^, Mg^2+^ and Phosphate. AFM **(A)** 2D image and **(B)** 3D image of 0.2 µM AkP. AFM **(C)** 2D image and **(D)** 3D image of 0.2 µM AkP with 0.5 mM Zn(NO_3_)_2_, 0.5 mM Mg(NO_3_)_2_ and 0.5 mM Na_2_HPO_4_. In the 2D images, white indicates sizes of 15 nm, while in the 3D images, the blue indicates sizes of 15 nm. The buffer used to make all experimental solutions was 50 mM HEPES (pH 7). The substrate solutions were filtered before being dropped on the mica in both experiments.

Lastly, we measured the activity of alkaline phosphatase using UV-Vis spectroscopy in the presence of pNPP with increasing added concentrations of zinc and magnesium ions. The reaction involving pNPP was used because the product, pNP, is UV active. We let the reaction proceed for 23 minutes to allow for aggregation and calculated the rate during the last three minutes. The results are shown in **Figure S4**. Most relevant are the cases with 0.5 mM and 1 mM zinc and magnesium nitrate, which correspond to the FRET and DLS experiments. As can be seen, while there is a small decrease in the activity of AkP, the decrease is not statistically significant, meaning that aggregates formed at these concentrations are still catalytically active.

#### Hexokinase aggregates with Mg^2+^ and either ATP or ADP at low pH values

Hexokinase (HK) from *Saccharomyces cerevisiae* was purchased in a lyophilized powder form from Millipore-Sigma. Its structure is shown in **Figure 6**. It phosphorylates D-Glucose (D-Glu) to glucose-6-phosphate (G6P) in the presence of magnesium chloride (MgCl_2_) and adenosine triphosphate (ATP) (Equation 3). It is a dimer with a weight of 110 kDa.

**Fig 6:**
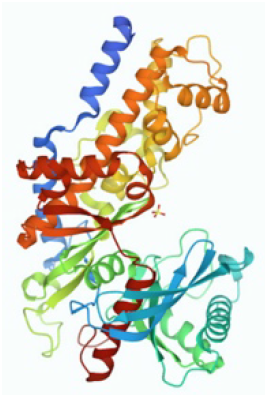
HK Structure. PDB structure of hexokinase (from *Saccharomyces cerevisiae)*.^32^

To begin, we examined the possible aggregation of HK in the presence of Mg^2+^ ion, which is required for activity,^33^ as well as cations and anions from the Hofmeister series. None of these ions had a discernible effect (**Figure S5**). Next, we tested the FRET efficiency with each of the species involved in HK catalysis, in the presence of the Mg^2+^ ion (**Figure S6**). As shown, neither D-Glu nor G6P has a significant effect on the aggregation (indeed, D-Glu causes fragmentation, see next section). However, both ADP and, especially ATP, in the presence of Mg^2+^, cause HK aggregation and an increase in the FRET efficiency signal (**Figure S6**).

We observed that adding 20 mM ATP or 20 mM ADP to the experimental solution caused a drop in the pH, despite the use of HEPES buffer, which has been noted in literature.^34^ Therefore, we tried two sets of experiments, one set in which we left the pH of the ATP or ADP solution as is and one in which we adjusted the pH of the ATP or ADP stock solutions to pH 7. The pH values for the stock solutions used to make the experimental solutions are found in **Table S2** of the SI and the pH of the resulting experimental solutions are found in **Figure 7** and in **Table S3** in the SI. **Table S3** contains the pH values for each of the experiments portrayed in **Figure 7**. The pH does not change dramatically as the experiment progresses, but as shown in **Figure 7**, lowering the pH significantly increases HK aggregation. The signals with HK; HK, Mg^2+^; HK, Mg^2+^, ATP; and HK, Mg^2+^, ADP all show higher levels of aggregation when the pH is left unadjusted, and the values are close to 4. When the pH is adjusted to 7, the levels of aggregation significantly drop. As with AkP, we used DLS to confirm the results we saw with FRET (**Figure S7**). Note the pH values were not controlled for these DLS experiments. The results from the DLS again suggests that the increase in FRET efficiency stems from the aggregation of the HK units, and not from the dissociation and recombination of monomeric subunits. It is worth noting that the pH remains at 7 in the FRET, DLS and AFM experiments involving AkP.

**Fig. 7.**
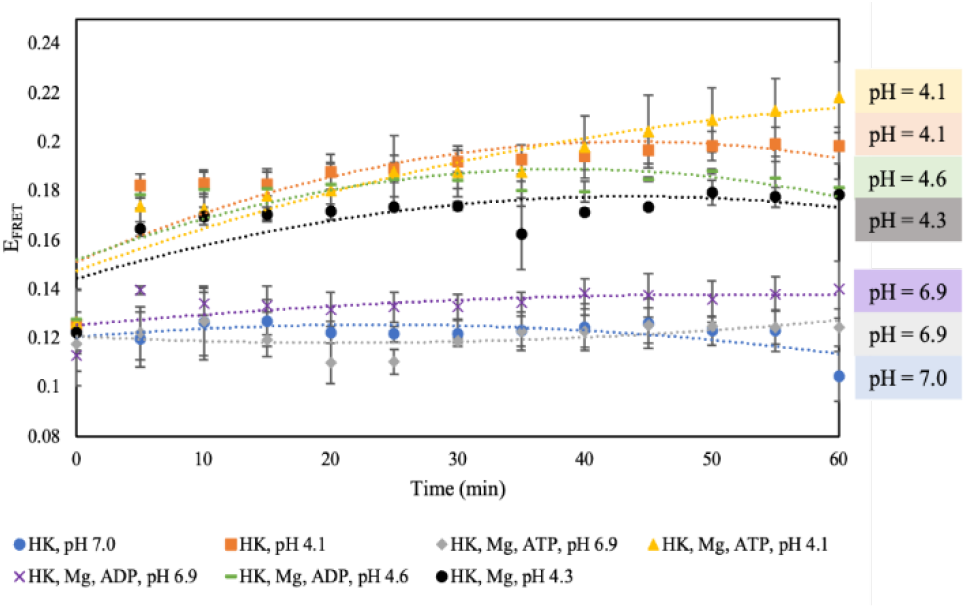
FRET efficiencies for HK at different pH values. FRET efficiency of 0.2 µM HK (pH 7.0; blue circles); 0.2 µM HK (pH 4.1; orange squares); 0.2 µM HK, 40 mM MgCl_2_, 20 mM ATP (pH 6.9; gray diamonds); 0.2 µM HK, 40 mM MgCl_2_, 20 mM ATP (pH 4.1; yellow triangles); 0.2 µM HK, 40 mM MgCl_2_, 20 mM ADP (pH 6.9; purple crosses); 0.2 µM HK, 40 mM MgCl_2_, 20 mM ADP (pH 4.6; green dashes); 0.2 µM HK, 40 mM MgCl_2_ (pH 4.3; black circles). The buffer used to make all experimental solutions was 50 mM HEPES (pH 7). The dashed lines are to guide the eye. The error bars represent the standard deviation from two trials.

Finally, we investigated the effect of aggregation on HK activity. We use a coupled assay with glucose 6-phosphate dehydrogenase (G6PDH) and ß-Nictoinamide Adenine Dinucleotide Phosphate (ß-NADP) to measure the activity of hexokinase with D-Glu, MgCl_2_ and ATP (Equations 3-4). We measured the rate of the reaction after 60 minutes and calculated the rate during the last three minutes. The results are seen in **Figure 8**. While some Mg^2+^ ion is required for HK optimal activity, the results for 20 mM and 40 mM Mg^2+^ appear to suggest that aggregation may result in an increase in the catalytic activity of HK, although the change is not statistically significant.

**Fig. 8.**
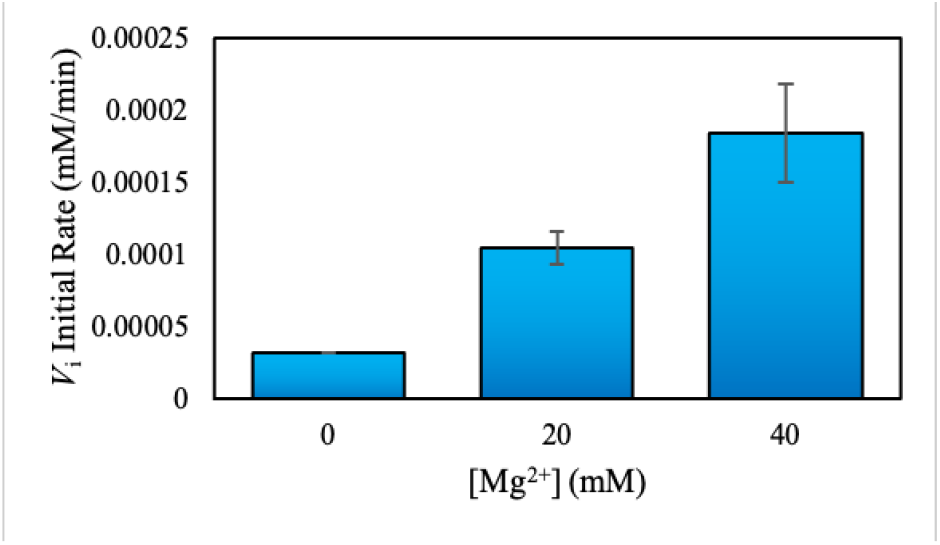
Activity of HK upon catalysis. Activity of 0.2 µM hexokinase between 57-60 min with 20 mM Glu, 20 mM ATP, 2.5 mM NADP^+^ and 10 units of G6PDH while increasing the concentration of MgCl_2_ from 0 to 40 mM. The buffer used to make all experimental solutions was 50 mM HEPES (pH 7). The error bars represent the standard deviation from two trials. The activities are not statistically different from 0 mM Mg^2+^ (P > 0.05; see Materials and Methods).

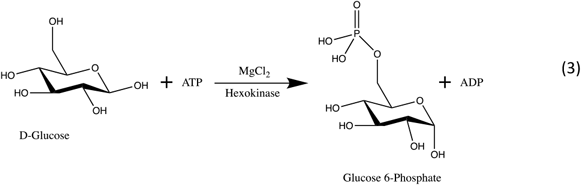

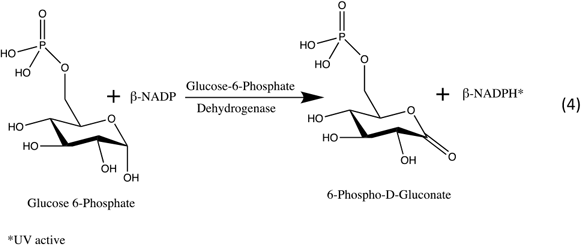

### Enzymes that Exhibit Fragmentation

#### Glucose Oxidase fragments upon the addition of D-Glucose

Glucose Oxidase (GOx) from *Aspergillus niger* was purchased from Millipore Sigma. Its structure is shown in **Figure 9**. This enzyme catalyzes the oxidation of D-Glucose (D-Glu) to form gluconic acid and hydrogen peroxide (Equation 5). It is a dimer and has a molecular weight of 160kDa.

**Fig 9:**
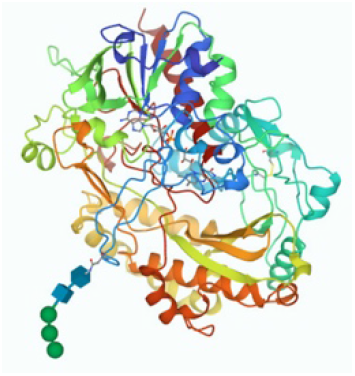
GOx Structure. PDB structure of glucose oxidase (*Aspergillus niger)*.^35^

Since glucose oxidase is not a metalloenzyme^36^, we did not investigate its aggregation behavior upon the addition of metal ions. Instead, using FRET, we examined the fragmentation of glucose oxidase in the presence of its substrate D-Glucose, the non-substrate enantiomer L-Glucose (L-Glu), the combined products of the reaction, as well as each product individually. (**Figure 10)**.

**Fig. 10.**
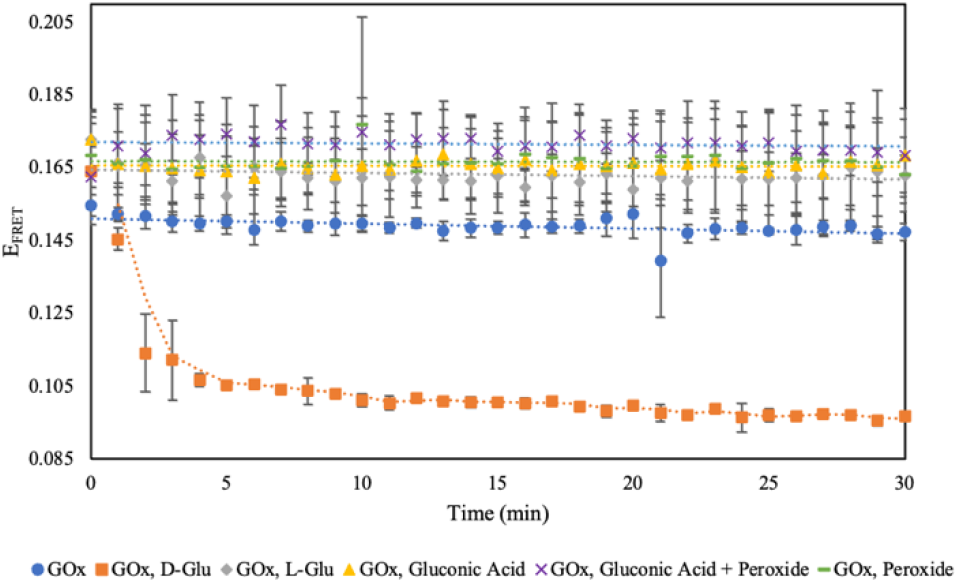
FRET efficiencies for GOx during catalysis. FRET efficiency for 0.2 µM GOx (blue circles); 0.2 µM GOx, 1 mM D-Glu (orange squares), 0.2 µM GOx,1 mM L-Glu (gray diamonds); 0.2 µM GOx, 1 mM gluconic acid (yellow triangles); 0.2 µM GOx, 1 mM gluconic acid and 1 mM hydrogen peroxide (purple crosses); 0.2 µM GOx, 1 mM hydrogen peroxide (green dashes). The buffer used to make all experimental solutions was 50 mM MES (pH 6). The dashed lines are to guide the eye. The error bars represent the standard deviation from three trials.

We found that the enzyme fragments in the presence of its substrate D-Glucose, but not L-Glucose. Enzymes typically have multimeric structures formed from polypeptide subunits. In principle, they can aggregate or fragment to larger or smaller multimeric structures. Therefore, we postulate that this fragmentation is due to the dissociation of the GOx enzyme into its free subunits.^28,37^ GOx does not fragment in the presence of the reaction products, gluconic acid and hydrogen peroxide. Furthermore, we investigated what happens if D-Glucose is not present in the reaction mixture initially but is produced gradually over time. To do so, we added another enzyme, invertase, to the solution. Invertase (Inv, also known as sucrase) converts sucrose to glucose and fructose as shown in Equation 6. Invertase from baker’s yeast (S. cerevisiae) was purchased from Millipore Sigma. Its structure is shown in **Figure 11**.

**Fig 11:**
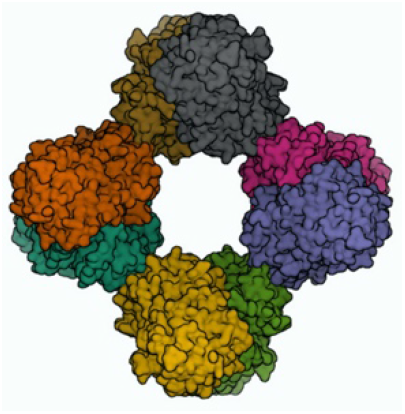
Inv Structure. PDB structure of invertase (from *Saccharomyces cerevisiae)*.^38^

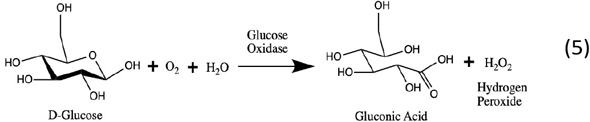

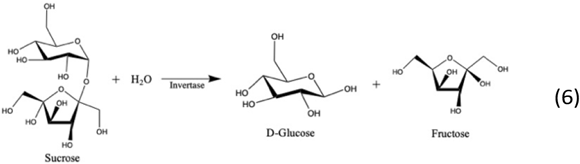

In this experiment, along with glucose oxidase tagged with the two dyes (AF488 and AF532), we added untagged 0.1 µM invertase. We tested the FRET efficiency of this solution by itself and after adding sucrose. We expected a time delay with a gradual onset of glucose oxidase fragmentation since invertase slowly converts sucrose to D-Glucose. The results are shown in **Figure 12**.

**Fig. 12.**
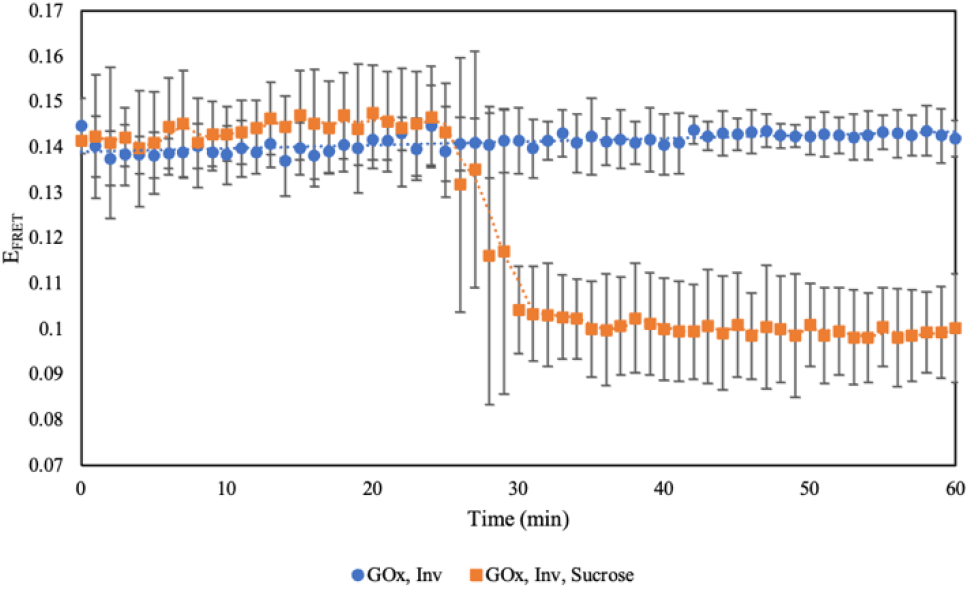
FRET efficiencies for GOx with Inv and Sucrose. FRET efficiency for 0.2 µM tagged GOx, 0.1 µM untagged Inv (blue circles); 0.2 µM GOx, 0.1 µM Inv, 1 mM sucrose (orange squares). the buffer used to make all experimental solutions was 50 mM MES (pH 6). The dashed lines are to guide The eye. the error bars represent the standard deviation from three trials.

As shown in **Figure 12**, we saw that, instead of fragmenting gradually, there was a sudden drop in the FRET efficiency which suggests that a minimum concentration of D-Glucose is required for GOx fragmentation. Based on preliminary calculations using the enzyme activity, we estimated that ∼0.3 mM D-Glucose was produced in 40 minutes after the addition of invertase and sucrose to the solution of glucose oxidase. These calculations are shown in the SI. We also performed concentration dependent experiments using FRET to obtain the concentration of D-Glucose at which the enzyme starts fragmenting. These results are summarized in **Figure S8**. They show that fragmentation starts after the addition of ∼0.3 mM D-Glucose which is similar to the value obtained in the calculations. Note that the physiological concentration of D-Glucose is 4.4-6.6 mM^39,40^, which is well above this concentration.

#### Hexokinase fragments in the presence of D-Glucose

We assessed the fragmentation behavior of hexokinase with both D- and the non-substrate enantiomer, L-Glucose. We found that D-Glucose by itself or D-Glucose in combination with ATP and MgCl_2_ causes a fragmentation as opposed to L-Glucose. These results are displayed in **Figure 13**.

**Fig. 13.**
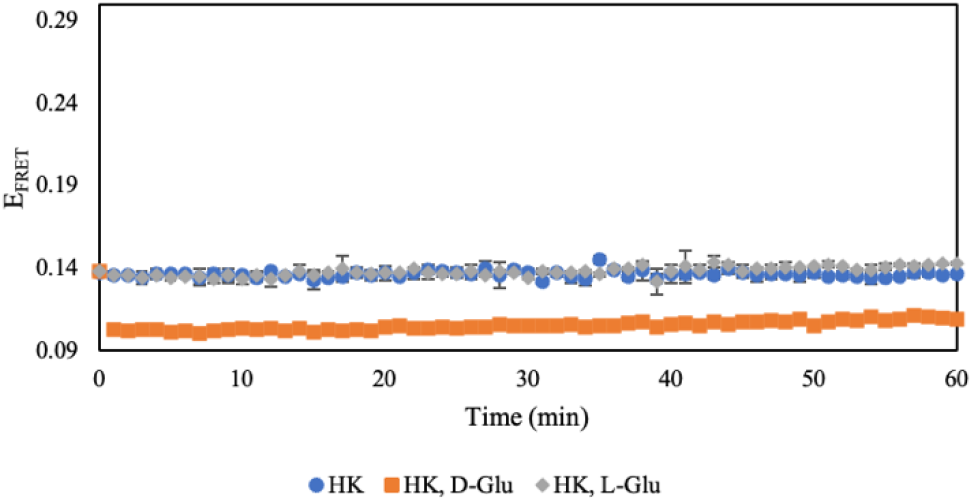
FRET efficiencies of HK with D- and L-Glu. FRET efficiency of 0.2 µM HK (blue circles); 0.2 µM HK, 20 mM D-Glu (orange squares); 0.2 µM HK, 20 mM L-Glu (gray diamonds). The buffer used to make all experimental solutions was 50 mM HEPES (pH 7). The error bars represent the standard deviation from two trials.

#### Alkaline Phosphatase does not fragment after the addition of D-Glucose

Based on the results shown above, we wondered whether the fragmentation by D-Glucose was only specific to those enzymes that use D-Glucose as the substrate (namely HK and GOx). Therefore, we examined the behavior of AkP which does not catalyze the transformation of D-Glucose. As is obvious from **Figure S9**, the addition of either D- or L-Glucose has no effect on the aggregation behavior of AkP.

## Conclusion

In this study, we have identified compounds that cause aggregation and fragmentation in several model enzymes. It is particularly noteworthy that these additives are directly involved in catalysis by the respective enzymes. Specifically, we found that glucose oxidase and hexokinase fragment in the presence of D-Glucose but not L-Glucose, while hexokinase aggregates in the presence Mg^2+^ ion and either ATP or ADP at low pH. Alkaline phosphatase aggregates in the presence of Zn^2+^ ion and inorganic phosphate. The aggregation of hexokinase and alkaline phosphatase does not appear to attenuate their catalytic activity. The results are summarized in **Figure 14**.

**Figure 14:**
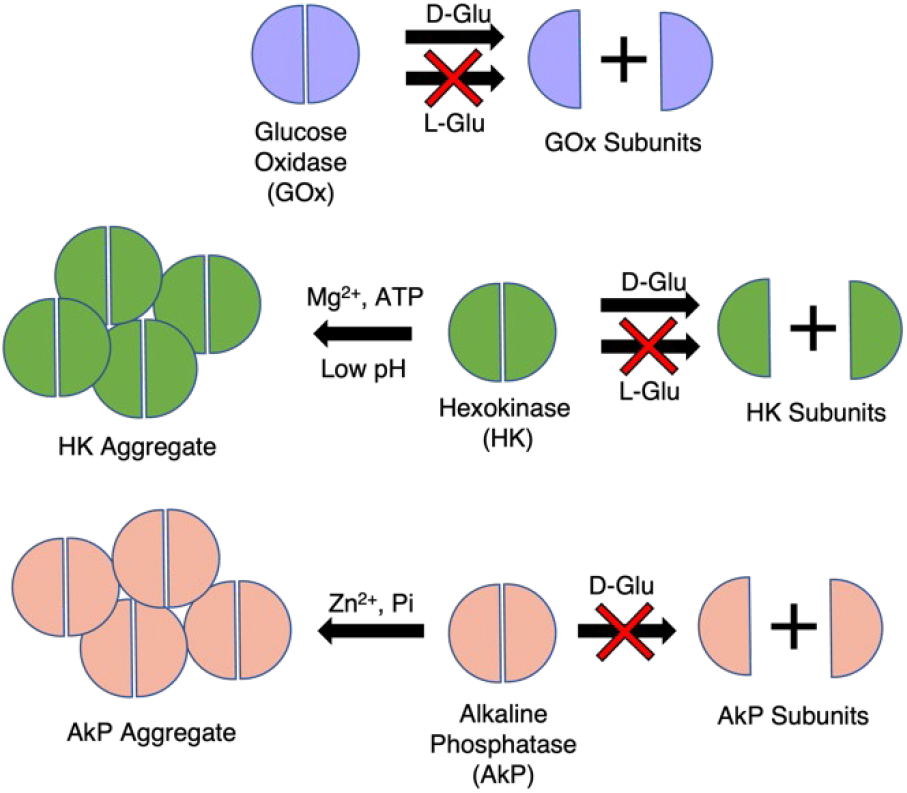
Schematic of enzyme aggregation and fragmentation. Glucose oxidase fragments into its subunits in the presence of D-Glu, but not L-Glu. Hexokinase also fragments to its subunits in the presence of D-Glu, but not L-Glu and aggregates in the presence of Mg^2+^ ion and ATP at low pH. Alkaline phosphatase aggregates in the presence of the Zn^2+^ ion and inorganic phosphate and does not fragment into its subunits in the presence of D-Glu

Our study underscores the dynamic nature of protein aggregates. It is clear that, for example, one should not *a priori* assume that the specific multimeric structure of the native enzyme is maintained during catalysis. Perhaps more interesting is that the work presented suggests pathways for different enzymes to associate or separate in the course of catalysis. Additionally, understanding the causes of enzyme dissociation or aggregation has multiple benefits in the industrial field.^41^ For example, the aggregation and fragmentation of multimeric enzymes must be taken into account in designing enzyme immobilization strategies. The mechanistic underpinnings of the processes are complex, and in this context, we note that a separate study has shown that modest concentrations of ATP can cause fragmentation and solubilization of disordered proteins.^42^

## Author Contributions

Conceptualization – KG, AB, JA, THL, AS

Supervision – JA, THL, AS

Funding acquisition – AS

Methodology – KG, AB, JK, SM, SG

Writing – original draft – KG, AB, AS

Writing – review & editing – KG, AB, JK, SG, SM, THL, AS

## Acknowledgements

The authors would like to thank Professor Christine Keating (Penn State) for allowing us the use of her fluorimeter for the FRET measurements. We also thank the rest of the Keating lab, especially Charlie Crowe, Saehyun Choi and Jennifer Miller for training us to use the fluorimeter, as well as accommodating us in their lab during the COVID-19 pandemic. We would also like to thank Kelly Collins from Brookhaven Instruments Corporation for her guidance and assistance in using the DLS. Finally, we thank Professor Henry Hess (Columbia) for helpful discussions. We gratefully acknowledge funding of the research by the Air Force Office of Scientific Research FA9550-20-1-0393 (AS).

